# Sharp wave-associated activity patterns of olfactory cortical neurons in the mouse piriform cortex

**DOI:** 10.1101/329185

**Authors:** Kazuki Katori, Hiroyuki Manabe, Ai Nakashima, Eer Dunfu, Takuya Sasaki, Yuji Ikegaya, Haruki Takeuchi

## Abstract

The olfactory piriform cortex is thought to participate in olfactory associative memory. Like the hippocampus, which is essential for episodic memory, it belongs to an evolutionally conserved paleocortex and comprises a three-layered cortical structure. During slow-wave sleep, the olfactory piriform cortex becomes less responsive to external odor stimuli and instead displays sharp wave (SPW) activity similar to that observed in the hippocampus. Neural activity patterns during hippocampal SPW have been intensively studied in terms of memory consolidation; however, little is known about the activity patterns of olfactory cortical neurons during olfactory cortex sharp waves (OC-SPWs). In this study, we recorded multi-unit neural activities in the anterior piriform cortex in urethane-anesthetized mice. We found that the activity patterns of olfactory cortical neurons during OC-SPWs were non-randomly organized. Individual olfactory cortical neurons varied in the timings of their peak firing rates during OC-SPW events. Moreover, specific pairs of olfactory cortical neurons were more frequently activated together than expected by chance. On the basis of these observations, we speculate that coordinated activation of specific subsets of olfactory cortical neurons repeats during OC-SPWs, thereby facilitating synaptic plasticity underlying the consolidation of olfactory associative memories.

## INTRODUCTION

Memories are encoded by synaptic modification, induced by coordinated activities of subsets of neurons during waking behavior, and consolidated during subsequent sleep. Neural activity patterns related to a waking experience have been replayed spontaneously during sleep. This replay has been studied intensively in the hippocampus, which is involved in spatial and episodic memory formation. Previous lesion studies suggest that olfactory associative memory is not stored in the hippocampus but rather in the piriform cortex (PC) (Staubli *et al.*, 1987; Bunsey & Eichenbaum, 1996; Fortin *et al.*, 2002; Chapuis *et al.*, 2013). The PC is the largest subregion of the olfactory cortex (OC), and in rodents it is located in the ventrolateral region of the brain. The PC receives massive afferent inputs from the olfactory bulb (OB), which is the first relay station of the olfactory system. There are several similarities between the PC and the hippocampus. The PC is an evolutionally conserved three-layered paleocortex. Further, olfactory cortical neurons generate massive numbers of recurrent association fibers within the PC, which are reminiscent of those of CA3 pyramidal cells in the hippocampus. In addition to the anatomical similarities, the PC generates characteristic local field potentials called sharp waves (SPWs) during slow-wave sleep period (Manabe *et al.*, 2011). SPWs are large negative deflections characteristic of the hippocampus and are implicated in memory consolidation (Buzsaki, 2015). In the hippocampus, specific subsets of pyramidal cells that were active during a previous waking period are repeatedly reactivated during later SPWs. There are several reports showing that SPW-associated reactivation of specific cell assemblies promotes consolidation of recent acquired memories (Girardeau *et al.*, 2009; Sadowski *et al.*, 2016).

It has been shown that olfactory cortex (OC)-SPWs are self-organized events that occur independently of hippocampal SPWs (Manabe *et al.*, 2011). The OC-SPWs propagate to various nearby areas of the brain such as the olfactory tubercle and olfactory bulb (OB) (Narikiyo *et al.*, 2014). OC-SPWs are related to the state of the orbitofrontal cortex, which has reciprocal connections with the anterior part of the PC; OC-SPWs preferentially occur at early and late phases of the “up” state of the orbitofrontal cortex (Onisawa *et al.*, 2017). Therefore, OC-SPWs are thought to provide a means of communication with olfactory-related brain areas.

Neural activity patterns during hippocampal SPWs have been intensively studied. In the hippocampus, specific cell ensembles are repeatedly activated during SPWs (Lee & Wilson, 2002), which play a role in memory consolidation. However, little is known about the activity patterns of olfactory cortical neurons during OC-SPWs. To gain insight into the roles of OC-SPWs, in the present study we acquired and analyzed multi-unit recordings of the activity patterns of cortical neurons during OC-SPWs in urethane-anesthetized mice. Individual cortical neurons showed variability in the timing of peak firing during OC-SPWs. Moreover, some neuron pairs were significantly more frequently recruited to OC-SPWs than chance would predict. These results imply that cortical neurons are activated in a coordinated manner during OC-SPWs.

## MATERIALS AND METHODS

### Animals

A total of 6 mice (C57BL/6; >P56, >25 g), including both male and female mice, were used in this experiment. All experimental procedures were performed with the approval of the animal experiment ethics committee at the University of Tokyo and according to the University of Tokyo guidelines for the care and use of laboratory animals.

### Surgery for tetrode implantation

The mice were anesthetized with isoflurane gas (1–1.5%), and an electrode assembly (microdrive), holding 5 simultaneously moving tetrodes and constructed using a 3-D printer, was stereotaxically implanted above the left piriform cortex (1.4 mm anterior and 1.7 mm lateral to the bregma, tilted 10°). The tip of the tetrode bundle was lowered to the cortical surface, and the tetrodes were inserted 3.5 mm into the brain at the end of surgery. Each tetrode was made of four twisted 17-μm polyimide-coated platinum-iridium (90/10%) wires (California Fine Wire). The electrode tips were plated with platinum to lower their impedances to 150–300 kΩ at 1 kHz. The tips of the tetrodes were coated with DiI (80 mg/ml, diluted with acetone). Two screws were threaded into the bone above the cerebellum for grounding. These electrodes were connected to an electrical interface board (Neuralynx) on the microdrive. After the microdrive was implanted above the piriform cortex, it was fixed in place using LOCTITE 454 (Henkel).

### Histology

After the recordings were completed, the mice were perfused transcardially with ice-cold 4% paraformaldehyde (PFA) in phosphate buffer solution (PBS). To aid reconstruction of the electrode tracks, the electrodes were withdrawn 1–2 hours after perfusion. After perfusion, the whole brains were post-fixed with 4% PFA in PBS overnight at 4°C. Coronal sections (100–200 μm) of the brain were made with a vibratome (Dosaka, Japan) and mounted on slides. The recording sites were then identified by tracking the fluorescence of DiI.

### Electrophysiological recordings in anesthetized mice

The tetrodes were simultaneously moved every day using a screw (Nogatadenki) until the tetrode tips reached layer II or III of the PC. After the tips reached layer II or III of the PC, the mice were anesthetized deeply with an intraperitoneal injection of urethane (1 g/kg body weight). For each mouse, recording was carried out for 30–60 minutes. Local field potential (LFP) recordings were sampled and digitized at 2 kHz and filtered with a bandpass of 0.1–500 Hz using a Cereplex direct recording system (Blackrock). Unit activity was amplified and band-pass filtered at 600 Hz to 6 kHz. Spike waveforms above a trigger threshold (60 μV) were time-stamped and recorded at 30 kHz for 1.6 msec each.

### Spike sorting

Spike sorting was performed offline using the graphical cluster-cutting software MClust (Redish, 2009). Clustering was performed manually in 2D projections of the multidimensional parameter space (waveform amplitudes, peak-to-trough amplitude differences, waveform energies, and the first principal component of waveform, each measured on the four channels of each tetrode). Clusters that yielded inter-spike intervals of less than 2 msec for more than 1% of the samples were excluded from the analysis.

### Identification of OC-SPWs

OC-SPWs were characterized as large sharp negative LFPs with a duration less than 200 msec. To identify OC-SPWs, the LFP from each tetrode was down-sampled to 100 Hz and bandpass filtered at 2–20 Hz. The baseline mean and standard deviation (SD) were calculated, and the threshold for SPW detection was set to 4 SDs above the mean. The time span of each OC-SPW was defined as the time of the amplitude peak ± 100 msec.

### Statistical analysis

All statistical analyses were performed with MATLAB software (Mathworks).

The probability of recruitment for each cell was defined as the fraction of OC-SPW events in which it fired at least once.

A cross-correlogram was created by calculating the number of interspike interval events of a cell pair within 10 msec bins. All recorded spikes were used in this analysis. The peak of the cross-correlogram was defined as the largest bin value. The peak was considered statistically significant if it exceeded the largest peak calculated from 100 surrogate datasets. Linear regression was calculated using the “fitlm” function in MATLAB.

Unless otherwise stated, all data are presented as mean ± s.e.m.

## RESULTS

### Individual cortical neurons in the piriform cortex showed variability in the timing of peak firing during OC-SPWs

To analyze activity patterns of cortical neurons during OC-SPWs, we implanted tetrodes into layer II and III of the anterior part of the PC where cell bodies of cortical neurons are located (Fig. 1A). To confirm recording sites, tetrodes were coated with fluorescent dye, DiI. (Figs. 1B and 1C). Mice were anesthetized with urethane to achieve stable long-term recordings (Manabe *et al.*, 2011). Simultaneous recordings of LFPs and action potentials revealed that a subset of cortical neurons showed increased spike activity during OC-SPWs (Figs. 1D and 2A). Individual cells showed variability in the timing of peak firing relative to the peak of the OC-SPW (Fig. 2B shows three examples). Some cortical neurons showed the highest spike activity before the peaks of OC-SPWs, whereas others showed it after the peaks (Fig. 2B). Then, z-scored firing rate histograms of all neurons recorded were averaged (Fig. 2C). Consistent with a previous report (Manabe *et al.*, 2011), we found a clear peak timed slightly before the OC-SPW peak, indicating that many neurons fired in the descending phase of OC-SPWs. Finally, we sorted the z-scored firing rates of cortical neurons by the timing of the peak firing rates during the OC-SPWs (Fig. 2D). The majority of cortical neurons fired in the descending phase of OC-SPWs. The timing of the peak firing rate for 26 of 65 cells was between −20 msec and 0 msec. This result suggests that each cortical neuron had a unique timing of peak firing during the OC-SPWs.

**Figure 1.**
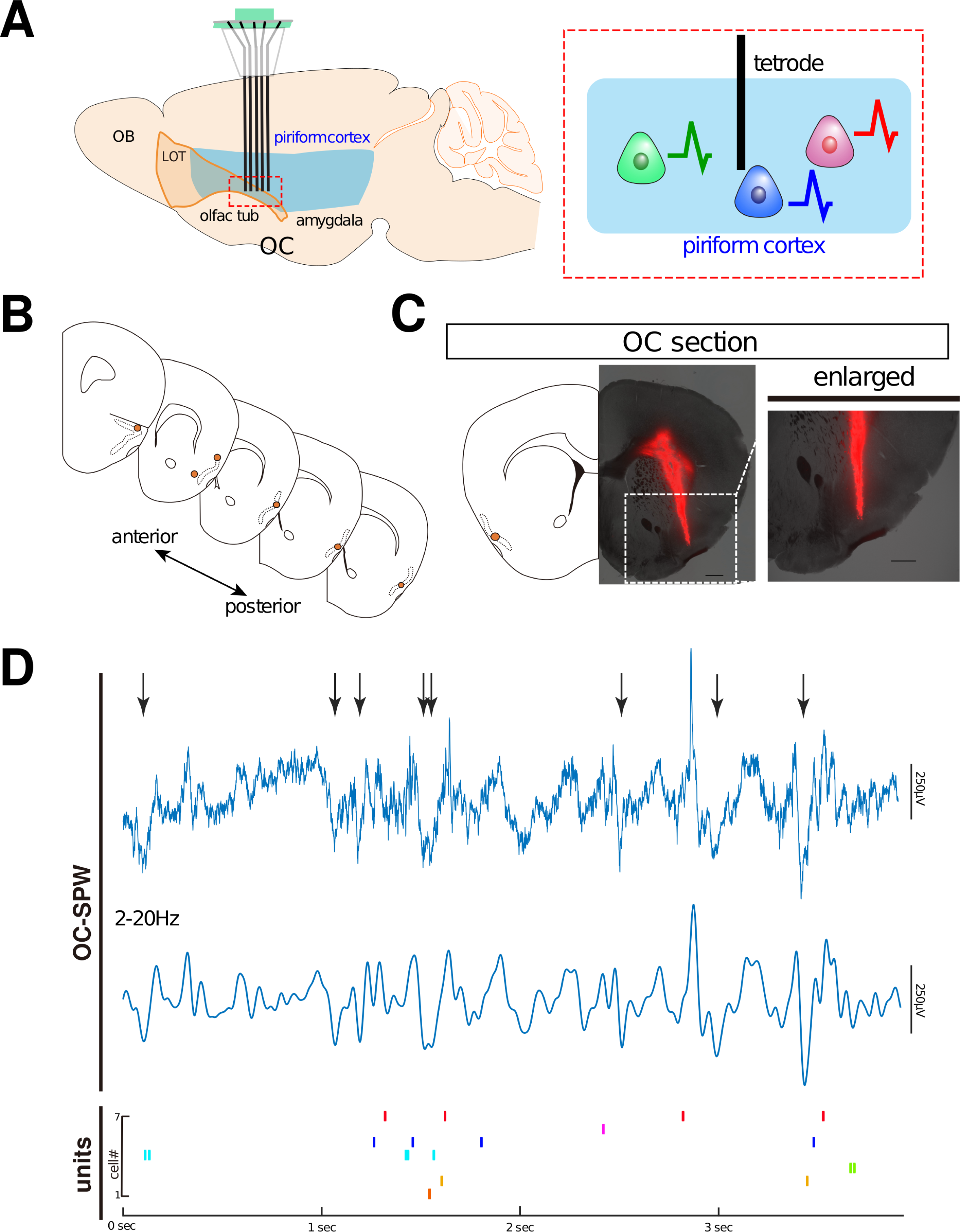
OC-SPWs occur in urethane-anesthetized mice. (A) Schematic illustration of recording methods. Tetrodes were implanted in the anterior part of the piriform cortex (PC; left). (B) Examples of tetrode locations are shown on schematic drawings of the brain (orange dots). The piriform cortex is shown by dotted line. (C) A representative coronal brain section showing the recording site in the PC. An enlarged view of the boxed area was shown on the right. DiI-coated tetrodes were used to determine the location of recording sites. Scale bar, 500 μm. (D) Simultaneous recordings of local field potentials (LFPs), band-pass filtered (2-20 Hz) trace, and single unit activities of cortical neurons. Arrows indicate olfactory cortex sharp waves (OC-SPWs). The raster plots represent unit recordings from 7 isolated PC neurons (bottom). OB, olfactory bulb; LOT, lateral olfactory tract; OC, olfactory cortex.

**Figure 2.**
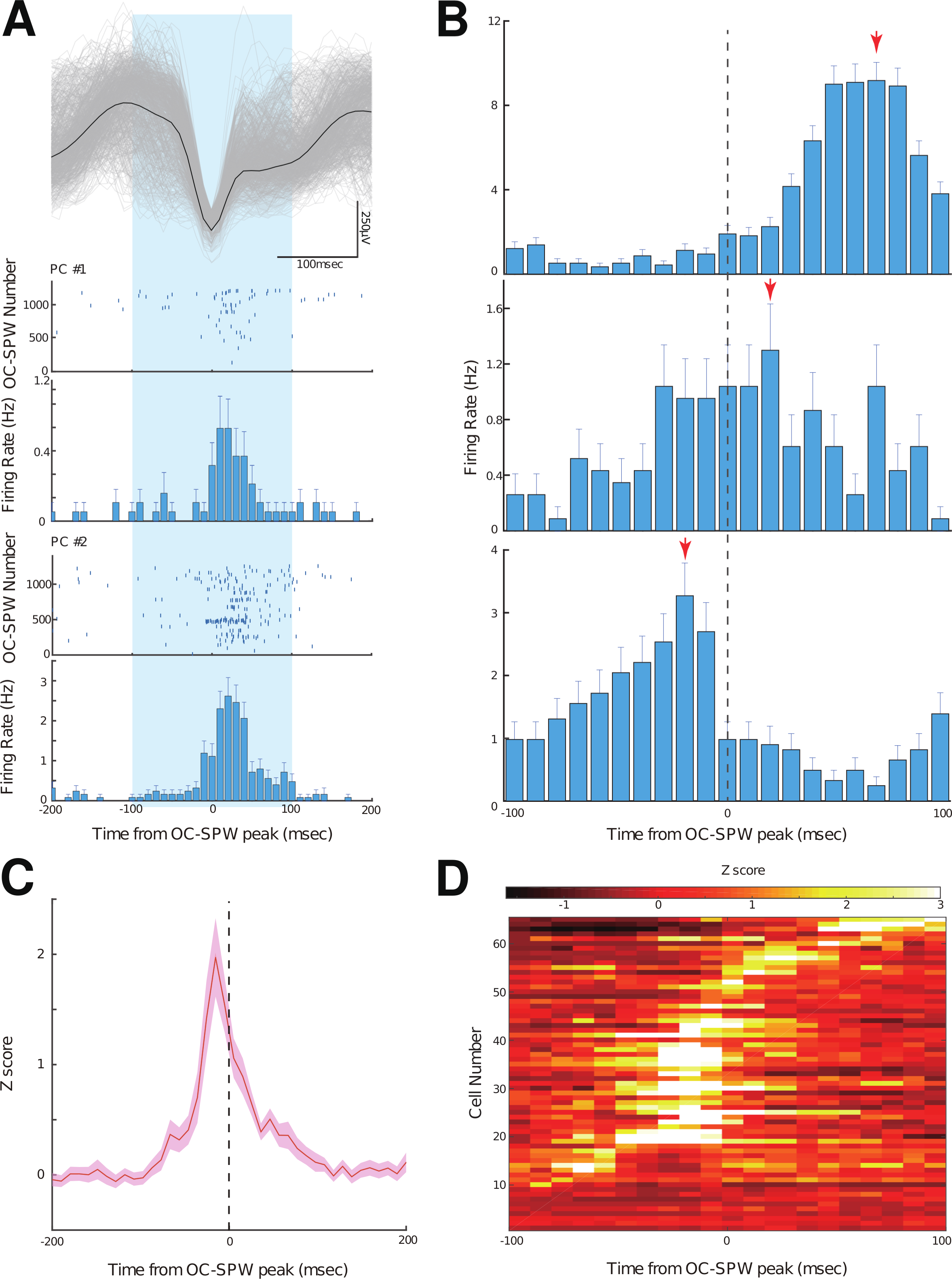
Individual cortical neurons in the anterior part of the piriform cortex showed variability in the timing of peak firing during olfactory cortex sharp waves (OC-SPWs) (A) Spike activities of cortical neurons during OC-SPWs. Top: Superimposition of all recorded OC-SPWs (n = 1261), aligned by the minima of the waveforms (gray lines). The black trace is the averaged waveform. Bottom: Raster plots and firing rate histograms (10 msec bins) of spike discharges from two isolated cortical neurons. Each row of the raster plots corresponds to one OC-SPW event. The time window is ± 100 msec, centered on the peak of the OC-SPW. (B) Firing rate histograms for 3 isolated cortical neurons during the OC-SPWs. Arrows indicate the timing of peak firing rates during the OC-SPWs. (C) Averaged Z-scored firing rate of all the cells recorded. The dotted line at the center indicates the timing of the peak of the OC-SPW. We used mean firing rates and SD values calculated from the entire recording period. (D) Temporal firing profiles of the cells during OC-SPWs, generated by the timing of the peak firing rates during the OC-SPWs.

### Repeated discharge of specific neuron pairs during OC-SPWs

Each OC-SPW recruited the discharge of a unique subset of cortical neurons (Fig. 3A). To quantify the degree of simultaneous recruitment of the discharge of cortical neurons, we introduced the surprise index *S*, which is calculated based on the statistical salience of the observed number of simultaneous-recruited events between a pair of neurons relative to chance. For further analyses, we used mice from which neural activities of more than 10 cells were recorded simultaneously (48 cells from 3 mice). If cell *i* and cell *j* are independently recruited to OC-SPWs, the probability *P*(*n*) that they exhibit *n* simultaneous-recruitment events for a total number of OC-SPW events *N* during recording is given by the Poisson equation:

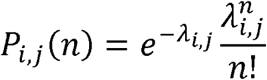

**Figure 3.**
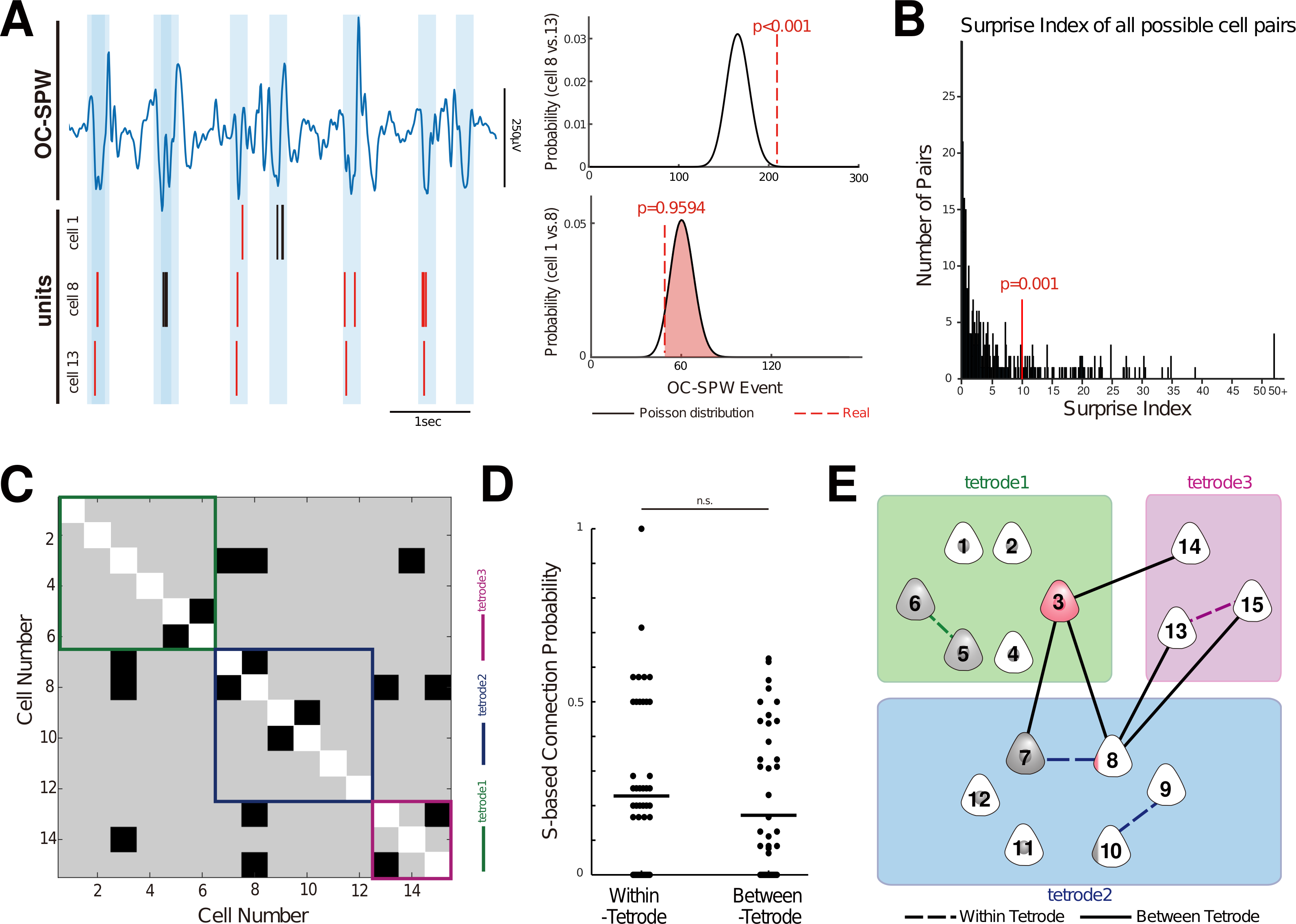
Coordinated activity patterns of cortical neurons during olfactory cortex sharp waves (OC-SPWs) (A) Trace of local field potentials (2-20 Hz) and raster plots of spike discharges of simultaneously recorded cells (cells #1, #8, and #13). The time windows of the OC-SPWs are shaded in blue. Action potentials associated with the OC-SPWs are shown in red (bottom). The plots on the right show a Poisson probability distribution of simultaneous recruitment events (black) and the number of real simultaneous recruitment events (red dotted line). The p value for each cell pair is indicated in red. For cells #8 and #13, the number of simultaneous recruitment events was significantly higher than chance. (B) Histogram of surprise indices. Neuron pairs showing a surprise index greater than 9.9658 (p < 0.001) were defined as simultaneously-recruited pairs. (n = 88 of 381 pairs) (C) Surprise index (S)-based connectivity matrix for simultaneously recorded cortical neurons (n = 15). The black and gray boxes indicate high (S > 9.9658) and low (S ≤ 9.9658) surprise indices, respectively. Cortical neurons that were recorded from the same tetrode (tetrode #1-3) are grouped and indicated in green, blue, and magenta, respectively. (D) The S-based connection probability (number of simultaneously-recruited pairs / number of possible pairs) was not significantly different between within-tetrode pairs and between-tetrode pairs (n = 48 cells from 3 animals, *p* **=** 0.0609, paired *t*-test). (E) A schematic diagram of connection patterns of cortical neurons. The dashed lines indicate connections between cells that were recorded from the same tetrode (within-tetrode), while solid lines indicate connections between tetrodes.

Where λ is the expected value of the number of simultaneous-recruitment events, i.e., *fi* ×*fj*×*N*, where *fi* and *fj* denote the probability of recruitment of cell *i* and cell *j*, respectively. When simultaneous-recruitment events are actually observed n times, the probability 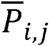 (rarity) is given as follows:

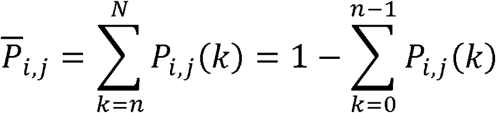

The surprise index *S* is then defined as follows: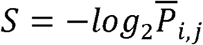.

Therefore, a pair of cortical neurons that fire frequently in the same OC-SPW events results in a high value of *S*. We calculated the surprise indices *S* for all possible pairs of cortical neurons and created an S-based connectivity matrix (Fig. 3B and 3C). Neuron pairs showing a surprise index greater than 9.9658 (p < 0.001) were defined as simultaneously-recruited pairs (n = 88 of 381 pairs). As shown in Fig. 3E, we found that the discharge of specific pairs of cortical neurons were recruited more frequently than chance would predict. For example, the discharge of cell #3 frequently occurred with those of cells #7, #8, and #14. Moreover, cell#8 was frequently recruited with cells #3, #7, #13, and #15. In contrast to cells #3 and #8, the discharge of cells #1, #2, #4, #11, and #12 did not occur with those of any other cells.

Simultaneously-recruited pairs were not found preferentially between recorded from the same tetrode and different tetrode (*p* = 0.0609, paired *t*-test) (Fig. 3D). This finding indicates that there is no spatial preference in the location of simultaneously-recruited pairs, which is consistent with the previous findings that association fibers of cortical neurons are broadly distributed in the PC, without any spatial preference (Franks *et al.*, 2011).

### Simultaneously-recruited pairs tended to fire at similar time

Finally, to examine the temporal relationship between pairs of cortical neurons, we constructed cross-correlograms for all possible pairs. Because of the small number of cortical neuron spikes during OC-SPWs, we incorporated all the spikes recorded, irrespective of OC-SPW recruitment. As shown in Fig. 4A, we found that some pairs showed a significant peak, whereas others showed flat distributions. Of a total of 381 pairs, 182 pairs showed significant peaks.

**Figure 4.**
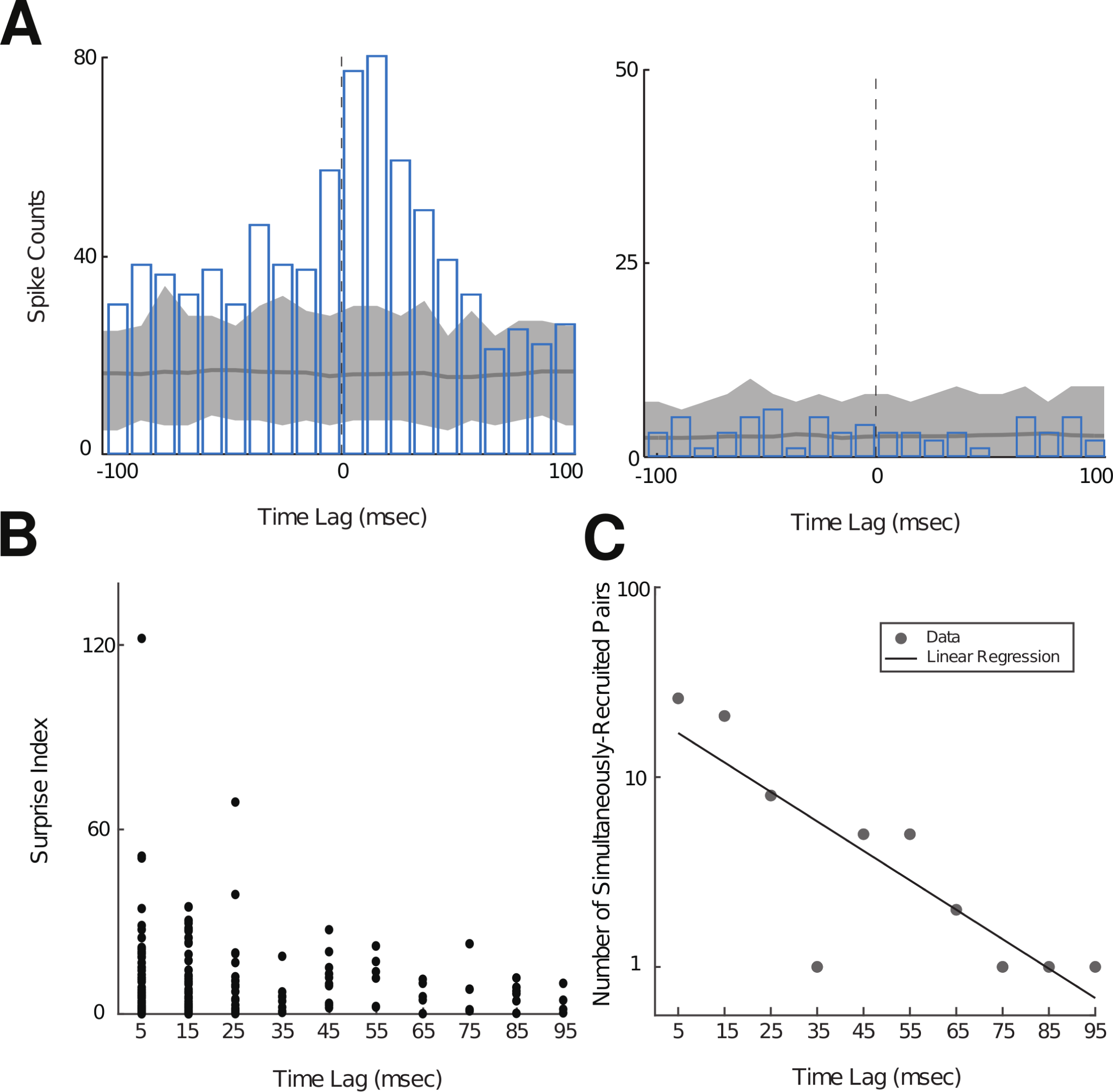
Relationship between surprise index and time lag of peak bin in cross-correlograms. (A) Cross-correlograms of cell pairs. Histograms show the number of spike counts for each 10 msec bin. Dark gray line shows averaged number of spike counts for each bin calculated from 100 sets of ISI-shuffled surrogates. The gray shaded area shows the range between the maximum number and minimum number of spike counts calculated from surrogates. Left: Cross-correlogram of a cell pair with a significant peak. Right: Cross-correlogram of a cell pair with no significant peak. (B) Relationship between the absolute time lag of peak bin and surprise index. <u>(R^2^ = 0.011, degrees of freedom = 180, *p* = 0.159, *F*-statistics)</u>. (C) The relationship between the absolute time lag of peak and the log number of simultaneously-recruited pairs. The black line indicates linear regression <u>(R^2^ = 0.65, degree of freedom = 8, *p* = 0.00485, *F*-statistics).</u>

We then examined the relationship between the peak time lag of the cell pairs and their surprise indices (Fig. 4B). No significant correlation was found (R^2^ = 0.011, degrees of freedom = 180, *p* = 0.159, *F*-statistics). However, we found a negative correlation between the peak time lag and the number of simultaneously-recruited pairs defined by the surprise index (R^2^ = 0.65, degree of freedom = 8, *p* = 0.00485, *F*-statistics) (Fig. 4C). These results suggest that the simultaneously-recruited pairs tend to fire at similar time.

## DISCUSSION

The PC is thought to be involved in both odor representation and odor associative memory (Staubli *et al.*, 1986; Barkai *et al.*, 1994; Thanos & Slotnick, 1997; Wilson & Stevenson, 2003). Olfactory information is initially represented as a spatially-ordered map of activated glomeruli in the OB. However, numerous studies have shown that the PC discards the spatially-ordered representation of odors; odor stimulus features are transformed into distributed patterns of neural activity in the PC (Stettler & Axel, 2009; Miura *et al.*, 2012; Bolding & Franks, 2017; Roland *et al.*, 2017). Although much effort has been paid to understand how odors are represented in the PC, little is known about the function of the PC in olfactory associative memory.

It is generally thought that memories are acquired during the waking period and consolidated during subsequent sleep periods. During slow-wave sleep period, the PC is less responsive to external odorants, and instead displays SPW activity. In the hippocampus, activity patterns from a previous waking period are repeatedly replayed during SPWs (Lee & Wilson, 2002). The SPW-associated neural activity is hypothesized to facilitate memory consolidation as well as enhance the link between hippocampal activity and neocortical activity (Ji & Wilson, 2007; Girardeau *et al.*, 2009; Peyrache *et al.*, 2009).

In the present study, we found coordinated activity of cortical neurons during OC-SPWs. Specific pairs of cortical neurons were frequently recruited to OC-SPWs. <u>However, we could not filter out firing of all of interneurons since electrical properties of interneurons in the PC have not been characterized. The coordinated activity pattern could be a result of combination of multiple phase-possessed cell types, as it has been shown different types of hippocampal interneurons show distinct phase possessions ((Sasaki *et al.*, 2014)). A previous report demonstrated that OC-SPWs preferentially occur during two distinct phases of the upstate of the orbitofrontal cortex (Onisawa *et al.*, 2017). In addition, we observed events where some OC-SPWs occurred close in timing. The OC-SPWs are likely to be classified into several types. It would be interesting to examine the relationship between activity patterns of cortical neurons and the types of OC-SPWs.</u>

It should be noted that no learning task was conducted in this study. Therefore, the coordinated activity patterns of cortical neurons observed in this study reflected an intrinsic property of the neural circuits in the PC. However, it has been shown that single-unit activity in the PC during slow-wave sleep is shaped by recent odor experience (Wilson, 2010). Moreover, slow-wave sleep imposed replay modulates both strength and precision of memory (Barnes & Wilson, 2014). In other sensory cortices, sequential activity patterns of neurons are broadly conserved between sensory stimuli and spontaneous activity during neocortical up-states. (Luczak *et al.*, 2007; Luczak *et al.*, 2009). It would be interesting to compare activity patterns of cortical neurons between sleep and wake periods doing some olfactory tasks.

Previous experiments have demonstrated that OC-SPWs generated in the PC can be observed in other olfactory-related brain areas, such as the olfactory tubercle (Narikiyo *et al.*, 2014) and olfactory bulb (OB) (Manabe *et al.*, 2011). It has been reported that pharmacological and electrical manipulation of top-down inputs from the PC to the OB during postprandial sleep affected plastic changes in intrabulbar circuits, by regulating the elimination of newly generated granule cells in the OB as a function of a prior waking experience (Yokoyama *et al.*, 2011; Komano-Inoue *et al.*, 2014). As granule cells play an important role in odor discrimination by modulating secondary olfactory neurons (Yokoi *et al.*, 1995; Shepherd *et al.*, 2007), re-organization of granule cells through top-down inputs from the PC during the postprandial sleep period may be important for olfactory information processing. In this scenario, OC-SPWs from the PC to the OB are possible candidates for such inter-regional information transfer. At any rate, further experiments, such as intervention in OC-SPWs, are necessary to address the function of OC-SPWs.

## ACKNOWLEDGEMENTS

This work was supported by the Yamazaki Spice Promotion Foundation, the Japan Science and Technology Agency, PRESTO, and the Japan Society for the Promotion of Science KAKENHI Grant Number, JPMJPR13AE, 16H06144, 16K14576, and 15K21029.

## AUTHOR CONTRIBUTIONS

KK, HM, AN, and HT conceived the experiments. KK, HM, AN, E, TS, YI, and HT wrote the manuscript. KK, AN, and HT performed the experiments and analyzed the data. All authors contributed to the writing and provided helpful comments. All authors read and approved the final manuscript.

## DATA ACCESSIBILITY

Data analyzed in this study is available from the corresponding author on reasonable request.

## ABBREVIATIONS

SPW: sharp wave
OC: olfactory cortex
OB: olfactory bulb
OE: olfactory epithelium
OR: olfactory receptor
OSN: olfactory sensory neuron

## REFERENCES

Barkai, E., Bergman, R.E., Horwitz, G. & Hasselmo, M.E. (1994) Modulation of associative memory function in a biophysical simulation of rat piriform cortex. Journal of neurophysiology, 72, 659–677.

Barnes, D.C. & Wilson, D.A. (2014) Slow-wave sleep-imposed replay modulates both strength and precision of memory. J Neurosci, 34, 5134–5142.

Bolding, K.A. & Franks, K.M. (2017) Complementary codes for odor identity and intensity in olfactory cortex. Elife, 6.

Bunsey, M. & Eichenbaum, H. (1996) Conservation of hippocampal memory function in rats and humans. Nature, 379, 255–257.

Buzsaki, G. (2015) Hippocampal sharp wave-ripple: A cognitive biomarker for episodic memory and planning. Hippocampus, 25, 1073–1188.

Chapuis, J., Cohen, Y., He, X., Zhang, Z., Jin, S., Xu, F. & Wilson, D.A. (2013) Lateral entorhinal modulation of piriform cortical activity and fine odor discrimination. J Neurosci, 33, 13449–13459.

Fortin, N.J., Agster, K.L. & Eichenbaum, H.B. (2002) Critical role of the hippocampus in memory for sequences of events. Nat Neurosci, 5, 458–462.

Franks, K.M., Russo, M.J., Sosulski, D.L., Mulligan, A.A., Siegelbaum, S.A. & Axel, R. (2011) Recurrent circuitry dynamically shapes the activation of piriform cortex. Neuron, 72, 49–56.

Girardeau, G., Benchenane, K., Wiener, S.I., Buzsaki, G. & Zugaro, M.B. (2009) Selective suppression of hippocampal ripples impairs spatial memory. Nat Neurosci, 12, 1222–1223.

Ji, D. & Wilson, M.A. (2007) Coordinated memory replay in the visual cortex and hippocampus during sleep. Nat Neurosci, 10, 100–107.

Komano-Inoue, S., Manabe, H., Ota, M., Kusumoto-Yoshida, I., Yokoyama, T.K., Mori, K. & Yamaguchi, M. (2014) Top-down inputs from the olfactory cortex in the postprandial period promote elimination of granule cells in the olfactory bulb. Eur J Neurosci, 40, 2724–2733.

Lee, A.K. & Wilson, M.A. (2002) Memory of sequential experience in the hippocampus during slow wave sleep. Neuron, 36, 1183–1194.

Luczak, A., Bartho, P. & Harris, K.D. (2009) Spontaneous events outline the realm of possible sensory responses in neocortical populations. Neuron, 62, 413–425.

Luczak, A., Bartho, P., Marguet, S.L., Buzsaki, G. & Harris, K.D. (2007) Sequential structure of neocortical spontaneous activity in vivo. Proc Natl Acad Sci U S A, 104, 347–352.

Manabe, H., Kusumoto-Yoshida, I., Ota, M. & Mori, K. (2011) Olfactory cortex generates synchronized top-down inputs to the olfactory bulb during slow-wave sleep. J Neurosci, 31, 8123–8133.

Miura, K., Mainen, Z.F. & Uchida, N. (2012) Odor representations in olfactory cortex: distributed rate coding and decorrelated population activity. Neuron, 74, 1087–1098.

Narikiyo, K., Manabe, H. & Mori, K. (2014) Sharp wave-associated synchronized inputs from the piriform cortex activate olfactory tubercle neurons during slow-wave sleep. J Neurophysiol, 111, 72–81.

Onisawa, N., Manabe, H. & Mori, K. (2017) Temporal coordination of olfactory cortex sharp-wave activity with up- and downstates in the orbitofrontal cortex during slow-wave sleep. J Neurophysiol, 117, 123–135.

Peyrache, A., Khamassi, M., Benchenane, K., Wiener, S.I. & Battaglia, F.P. (2009) Replay of rule-learning related neural patterns in the prefrontal cortex during sleep. Nat Neurosci, 12, 919–926.

Roland, B., Deneux, T., Franks, K.M., Bathellier, B. & Fleischmann, A. (2017) Odor identity coding by distributed ensembles of neurons in the mouse olfactory cortex. Elife, 6.

Sadowski, J.H., Jones, M.W. & Mellor, J.R. (2016) Sharp-Wave Ripples Orchestrate the Induction of Synaptic Plasticity during Reactivation of Place Cell Firing Patterns in the Hippocampus. Cell Rep, 14, 1916–1929.

Sasaki, T., Matsuki, N. & Ikegaya, Y. (2014) Interneuron firing precedes sequential activation of neuronal ensembles in hippocampal slices. Eur J Neurosci, 39, 2027–2036.

Shepherd, G.M., Chen, W.R., Willhite, D., Migliore, M. & Greer, C.A. (2007) The olfactory granule cell: from classical enigma to central role in olfactory processing. Brain Res Rev, 55, 373–382.

Staubli, U., Fraser, D., Kessler, M. & Lynch, G. (1986) Studies on retrograde and anterograde amnesia of olfactory memory after denervation of the hippocampus by entorhinal cortex lesions. Behav Neural Biol, 46, 432–444.

Staubli, U., Schottler, F. & Nejat-Bina, D. (1987) Role of dorsomedial thalamic nucleus and piriform cortex in processing olfactory information. Behav Brain Res, 25, 117–129.

Stettler, D.D. & Axel, R. (2009) Representations of odor in the piriform cortex. Neuron, 63, 854–864.

Thanos, P.K. & Slotnick, B.M. (1997) Short-term odor memory: effects of posterior transection of the lateral olfactory tract in the rat. Physiol Behav, 61, 903–906.

Wilson, D.A. (2010) Single-unit activity in piriform cortex during slow-wave state is shaped by recent odor experience. The Journal of neuroscience : the official journal of the Society for Neuroscience, 30, 1760–1765.

Wilson, D.A. & Stevenson, R.J. (2003) Olfactory perceptual learning: the critical role of memory in odor discrimination. Neuroscience and biobehavioral reviews, 27, 307–328.

Yokoi, M., Mori, K. & Nakanishi, S. (1995) Refinement of odor molecule tuning by dendrodendritic synaptic inhibition in the olfactory bulb. Proc Natl Acad Sci U S A, 92, 3371–3375.

Yokoyama, T.K., Mochimaru, D., Murata, K., Manabe, H., Kobayakawa, K., Kobayakawa, R., Sakano, H., Mori, K. & Yamaguchi, M. (2011) Elimination of adult-born neurons in the olfactory bulb is promoted during the postprandial period. Neuron, 71, 883–897.

